# Detection of dedifferentiated stem cells in *Drosophila* testis

**DOI:** 10.1101/2025.03.06.641800

**Authors:** Muhammed Burak Bener, Boris M. Slepchenko, Mayu Inaba

## Abstract

Tissue homeostasis relies on the stable maintenance of the stem cell pool throughout an organism’s lifespan. Dedifferentiation, a process in which partially or terminally differentiated cells revert to a stem cell state, has been observed in a wide range of stem cell systems, and it has been implicated in the mechanisms for stem cell maintenance. Dedifferentiated stem cells are morphologically indistinguishable from original stem cells, making them challenging to identify. Therefore, whether dedifferentiated stem cells have any distinguishable characteristics compared with original stem cells is poorly understood. The *Drosophila* testis provides a well-established model to study dedifferentiation. While our previous live imaging analyses have identified dedifferentiation events constantly occurring at steady state, existing genetic marking methods fail to detect most of the dedifferentiated stem cells and thus significantly underestimate the frequency of dedifferentiation events. Here, we established a genetic tool with improved sensitivity and used live imaging and mathematical modeling to evaluate the system. Our findings indicate that the specificity of lineage-specific promoters is critical for successfully identifying dedifferentiated stem cells.

## Introduction

Tissue homeostasis and repair rely on a stable stem cell pool, and deficiencies in stem cells lead to tissue degeneration, dysfunction and disease. Therefore, it is essential to maintain the stem cell pool throughout the lifespan of an organism. Dedifferentiation, where partially or terminally differentiated cells revert to the stem cell state, is one mechanism proposed to support stem cell maintenance across various tissues during both steady-state and regeneration conditions [1–9]. Given its role in stem cell maintenance, studying dedifferentiation and identifying dedifferentiated stem cells is essential. However, the underlying mechanisms of dedifferentiation remain poorly understood.

A key challenge in identifying dedifferentiated stem cells is that they are often morphologically and molecularly similar to the original stem cells [4]. Genetic marking, such as a lineage tracing technique, has been employed to distinguish dedifferentiated stem cells from the original stem cells [3–5, 10–14]. Genetic marking often relies on promoters active in differentiated cells, but not in stem cells, to induce recombination of reporter cassettes, resulting in the permanent marking of these cells. Ideally, recombination should remain completely silent in stem cells while being sufficiently high in targeted cells for accurate marking. However, this has been proven to be difficult because many cell-stage-specific markers exhibit overlapping expression across stages (reviewed in [2]), resulting in either leaky reporter activation in stem cells due to basal recombinase expression or insufficient reporter activation in differentiating cells, reducing the accuracy of tracing tools [15].

The *Drosophila* testis serves as an excellent model for studying *in vivo* stem cell maintenance [16]. Each testis contains 8 to 12 germline stem cells (GSCs) around a cluster of post-mitotic cells called the hub [17]. GSCs typically divide asymmetrically, producing one daughter cell that remains attached to the hub and retains its stem cell identity, while the other daughter cell is detached from the hub (called gonialblasts or GBs) and starts the differentiation process [18]. A GB undergoes four rounds of mitosis to form cysts of 2-, 4-, 8-, and 16-cell spermatogonia (SG) [19].

Dedifferentiation can be experimentally induced using forced differentiation of stem cells in *Drosophila* testis. When the differentiation factor, Bag of Marbles (Bam), is temporarily expressed, all GSCs start differentiation and physically leave the hub. After the treatment, cells detached from the hub start migrating back to the hub and regaining stem cell identity [1]. This phenomenon was first described by Brawley et al., in 2004 [1]. However, how frequently dedifferentiation occurs under steady-state conditions is still unclear.

Current genetic marking tools detect dedifferentiated GSCs at low frequencies [10, 11]. All previous tools utilized the bamGal4 driver to drive FLP recombinase (UAS-FLP) expression and permanently mark differentiating cells (Figure 1B-D). The low efficiency of marking is likely due to the delay of FLP expression, which may not reach the threshold for effective recombination until the 4-cell SG stage or later. Since GBs are the primary source of dedifferentiation [9], such delay significantly affects the marking efficiency. Consequently, these existing tools may fail to capture dedifferentiated GSCs across the fly’s lifespan fully [10, 11].

**Figure 1.**
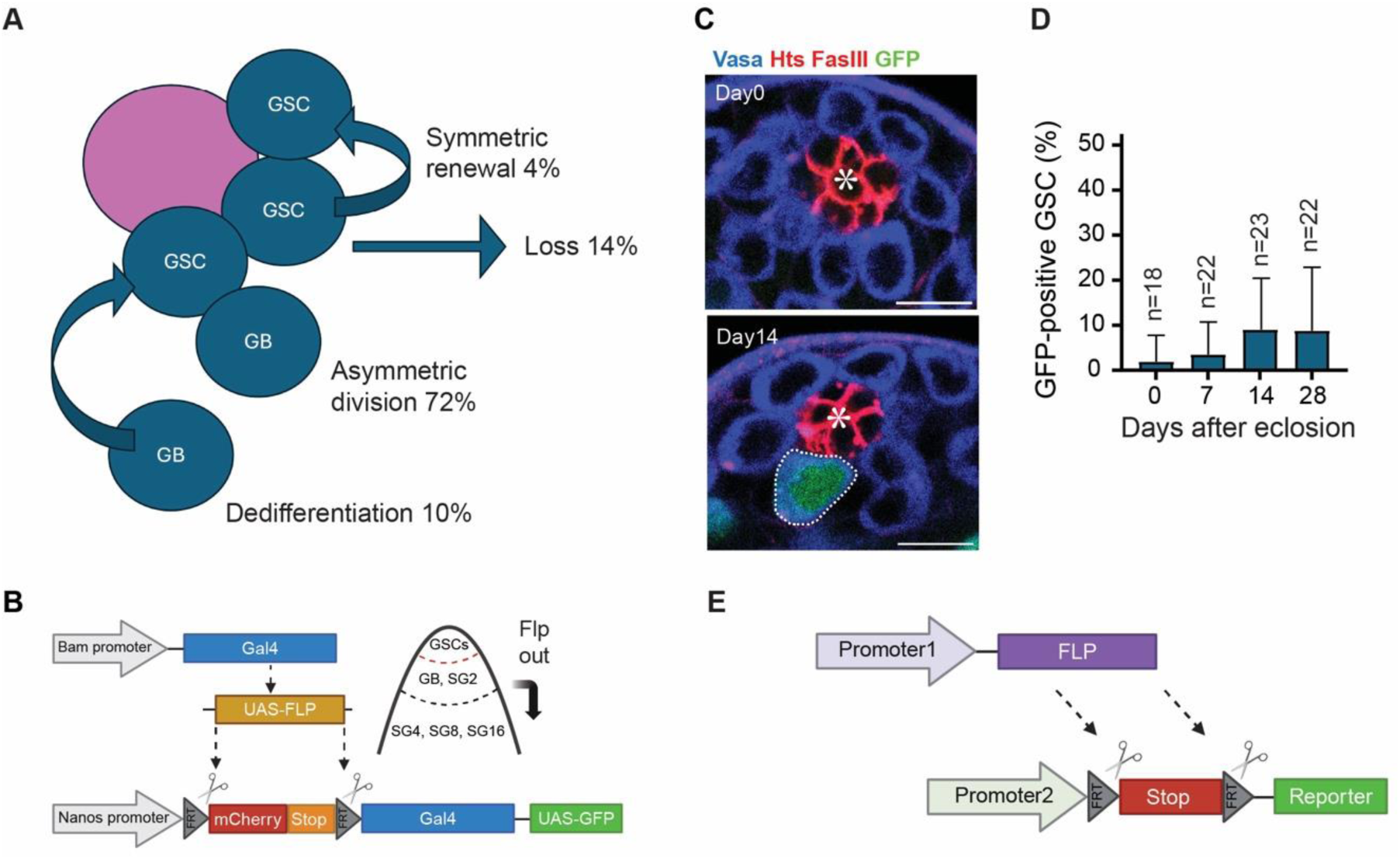
Improvement of genetic marking system for identifying dedifferentiated GSCs. **A)** A schematic diagram illustrating the estimated rates of asymmetric division, symmetric renewal, differentiation, and dedifferentiation of GSCs under steady-state conditions within the testicular niche shown in our previous study [9]. **B)** An illustration of the bamGal4-driven FLP lineage tracing system. The bamGal4 driver induces FLP expression in differentiating cells (4-cell and later-stage SGs), where FLP excises the mCherry-stop cassette. If cells with the removed stop cassette revert to the GSC state, they can be identified by the GFP expression under the control of nosGal4 driver. **C**) Representative immunofluorescence images showing the testicular niche in bamGal4-driven FLP lineage tracing system in the males at 0 and 14 days post-eclosion. The testes were stained with Vasa (blue), FasIII (red), and Hts (red) antibodies. GFP-positive GSCs (green) are encircled by white dotted lines. **D**) A graph showing the percentage of GFP-positive GSCs per testis at 0, 7, 14, and 28 days post-eclosion. **E)** An illustration of the lineage tracing system where FLP recombinase is directly expressed under a promoter and excises the FRT sites from the reporter cassette. The reporter gene is expressed under a separate promoter. All scale bars are 10 μm. Asterisks in the images indicate the approximate location of the hub. “n” indicates the number of testes scored.

In this study, we aimed to develop tools to improve efficiency. Specifically, we utilized FLP directly driven by the Bam promoter combined with FRT-flanked cassettes. We tested two FRT cassettes, each exhibiting a distinct threshold sensitivity to FLP dose. To evaluate the efficiency and specificity of this system, we conducted long-term live imaging. We found that the new FLP transgene has better efficiency, albeit with incomplete marking of dedifferentiated stem cells in the *Drosophila* testicular niche.

## Results

### Improvement of genetic marking system for identifying dedifferentiated GSCs

In our previous study, we investigated the mechanisms of GSC replenishment in the *Drosophila* testis niche under steady-state conditions using long-term live imaging (∼16 hours) [9]. Among all the events observed in the niche (∼6 total events per recording), 72% were asymmetric cell division (ACD), 14% were losses (symmetric differentiation or stem cell loss without division), 4% were symmetric renewal, and 10% were dedifferentiation (Figure 1A). Given this 10% dedifferentiation rate observed over 16-hour imaging period (∼0.6 dedifferentiation events per recording), we estimate that roughly ∼1 GSC will be supplied via dedifferentiation every day [9].

Previous studies have employed lineage-tracing tools to detect dedifferentiated GSCs [10, 11]. These systems rely on bamGal4-driven Flippase (FLP) expression that removes FRT-flanked stop codons located between the promoter and the downstream reporter coding sequence. When FLP-recombined cells dedifferentiate and become GSCs, these cells will be positive for LacZ or GFP [10, 11]. We first tested this previous system using nos-FRT-mCherry-stop-FRT-Gal4 cassette with UAS-GFP (hereafter referred to as nos>>GFP). On average, GFP-positive GSCs comprised only approximately 10% of the total GSC pool at 28 days post-eclosion (Figure 1B-D), similar to the previous report (∼14% shown in [11]). Considering the frequency observed in live imaging (1∼2/day [9]), this system might miss most dedifferentiation events. We speculate that this issue was likely due to delays in FLP expression when the Gal4/UAS binary system was used. Therefore, we sought to establish a new system.

To improve marking efficiency, we generated a transgenic fly to express FLPD5 (an FLP variant with enhanced activity [20]) directly driven by the Bam promoter so that we can bypass the Gal4-mediated UAS induction to minimize delays in inducing FLP recombinase activity (Figure 1E).

### Bam-FLPD5 causes spontaneous flipping events in GSCs, which obscures dedifferentiation marking

The newly established Bam-FLPD5 transgene showed a high frequency of GFP-positive cells after introducing the same nos>>GFP reporter cassette as in the previous system (Figure 2A-D). To evaluate whether the marked GFP-positive GSC frequency reflects the dedifferentiation rate, we conducted long-term live imaging of the niche to monitor GFP activation. Testes expressing both the Bam-FLPD5, nos>>GFP, and Vasa-mCherry (a germline cell marker) were imaged for approximately 12 hours using a z-stack to cover the entire niche. In 27 niches, each imaged for 12 hours, we observed eleven GSCs becoming GFP positive within the niche (Figure 2E, Movie S1), even though the Bam promoter used to drive FLP expression was expected to be active specifically in GB/SGs and not in GSCs. This is likely due to a leaky/nonspecific driver activity in GSCs, resulting in unintended flipping events. Within the same time period, the number of GB/SGs that became GFP-positive was twelve, indicating almost no difference in the frequency of flipping events in the GSC and GB/SG pools. Given the near-equal Bam-FLPD5 activity in GSC and GB/SG populations, we conclude that the majority of the GFP-positive GSCs were not generated by dedifferentiation events, and therefore this system is unsuitable for tracing dedifferentiated cells.

**Figure 2.**
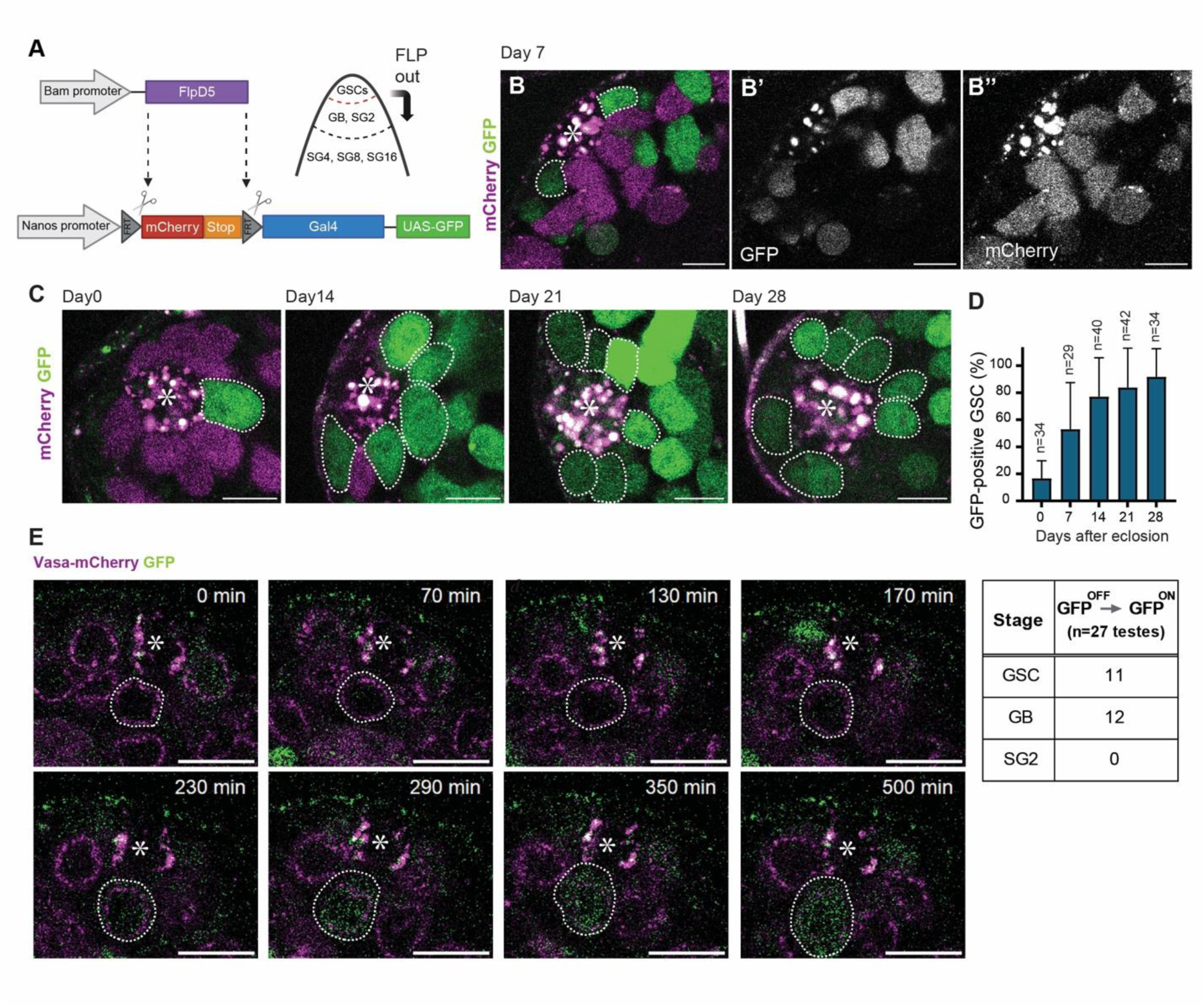
Bam-FLPD5 causes spontaneous flipping events in GSCs, which obscures dedifferentiation marking. **A)** An illustration of the Bam-FLPD5 lineage tracing system. Here, FLPD5 expression is driven directly under the Bam promoter sequence (-198 to transcription start site, TSS) [45]. The Bam-FLPD5 is crossed with nos-FRT-mCherry-stop-FRT-Gal4, UAS-GFP cassette (nos>>GFP) **B, C**) Representative images of live testes of indicated ages of the Bam-FLPD5-driven lineage-tracing system. GFP and mCherry channels are shown in separate panels in **B’** (for GFP) and **B’’** (for mCherry). **D)** A graph showing the scored percentage of GFP-positive GSCs per testis on days 0, 7, 14, 21, and 28 post-eclosion. **E)** Selected frames of time-lapse images from a 12-hour recording (interval:10min) of a niche showing the GFP-activation in a native GSC (corresponding to Movie S1). Germline cells are visualized by expression of Vasa-mCherry (magenta). White dotted lines encircle a GSC, becoming GFP-positive. Asterisks in the images indicate the approximate location of the hub. The table (right panel) shows the number of times GFP was observed to activate for each indicated cell stage (GSC, GB, and SG) during the 12-hour live imaging recording of 27 testes. All scale bars are 10 μm. Asterisks in the images indicate the approximate location of the hub. “n” indicates the number of testes scored.

### UAS-FRT-stop-FRT-mCD8-GFP cassette shows reduced spontaneous flipping rate in GSCs

We next strategized to overcome the issue of the nonspecific marking. Since the activity of Bam promoter in GSCs is expected to be weak [21, 22], Bam-FLPD5 likely yields low levels of FLP recombinase in the stem cells in the niche. On the other hand, the substrate constructs with FRT sites may differ in their sensitivity to FLP recombinase activity [15]. Therefore, we attempted to use an alternative reporter cassette with a higher threshold for FLP mediated recombination to minimize spontaneous GFP activation in GSCs, so that it responds linearly to FLP activity differences.

First, to test the recombination rates of different FRT cassettes, we compared the recombination frequency of nos>>GFP to an alternative UAS-FRT-stop-FRT-mCD8-GFP, together with nosGal4 (hereafter referred to as nos>>mCD8GFP) using heat-shock activatable promoter-FLP (hs-FLP). We performed a single 30-min heat-shock treatment at 37°C to induce FLP expression, then examined GFP positive GSC frequency after 24 hours [23, 24]. The nos>>mCD8GFP yielded a significantly lower percentage of GFP-positive GSCs than nos>>GFP indicating that nos>>mCD8GFP has a higher threshold to FLP and may be a good candidate for Bam-FLPD5 substrate for marking of dedifferentiated cells (Figure S2A-C).

Then, we examined the frequency of GFP-positive GSCs using a combination of Bam-FLPD5 and nos>>mCD8GFP and found that it indeed marks GSCs at a lower rate (Figure 3A-C) than the Bam-FLPD5, nos>>GFP system (Figure 2D). To assess the recombination rate of Bam-FLPD5 with nos>>mCD8GFP cassette in GSCs and in GB/SGs, we again employed a live imaging approach. During imaging, we observed frequent GFP activation in differentiating cells, including twelve GBs and four SGs, across 25 niches imaged for ∼12 hours each (Figure 3D, Movie S2). Notably, from these 25 recordings, we observed only one instance of GFP activation that occurred in GSCs without prior dedifferentiation events (Figure 3D).

**Figure 3.**
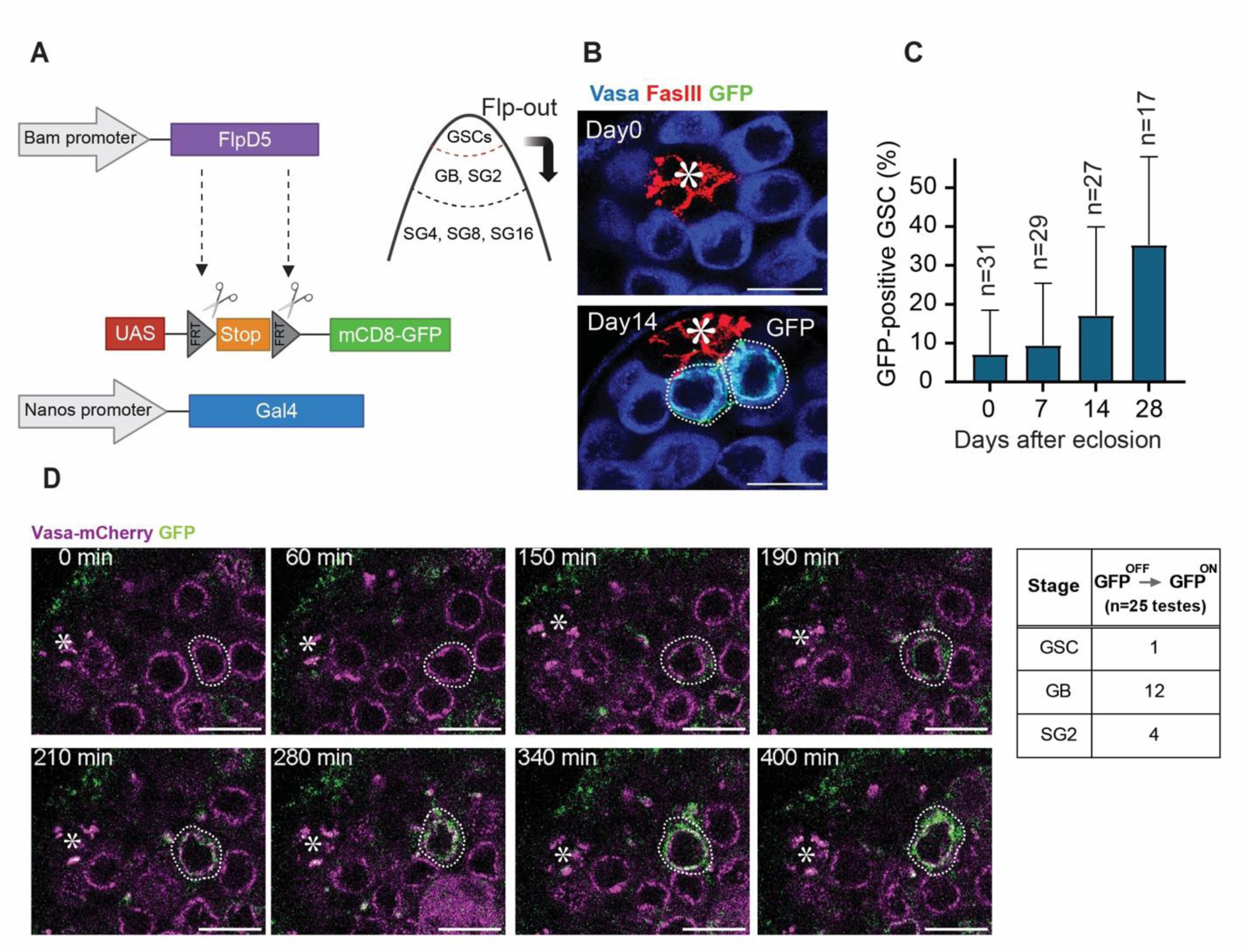
UAS-FRT-stop-FRT-mCD8-GFP cassette is efficiently flipped in GB/SGs but not in GSCs. **A)** An illustration of the Bam-FLPD5 lineage tracing system. Here, Bam-FLPD5 is combined with the UAS-FRT-stop-FRT-mCD8-GFP cassette and nosGal4 (nos>>mCD8GFP). FLPD5 removes the stop codon from this cassette. If cells once removed stop cassette in the past become GSCs, they can be identified by the GFP expression under the control of nosGal4 driver. **B)** Representative immunofluorescence images of the testicular niche of the Bam-FLPD5 marking system in the males at 0 and 14 days post-eclosion. The testes were stained with Vasa (blue) and FasIII (red) antibodies. GFP-positive GSCs (green) are outlined by white dotted lines. **C)** A graph showing the percentage of GFP-positive GSCs per testis at 0, 7, 14, and 28 days post-eclosion. **D)** Selected frames from a 12-hour time-lapse recording (interval:10 min) of a niche showing the GFP-activation in a GB (corresponding to Movie S2). Germline cells are visualized by expression of Vasa-mCherry (magenta). White dotted lines encircle a GB becoming GFP positive. The table (right panel) shows the number of times GFP was observed to activate for each indicated cell stage (GSC, GB, and SG) during 12-hour live imaging recording of 25 testes. All scale bars are 10 μm. Asterisks in the images indicate the approximate location of the hub. “n” indicates the number of testes scored.

The observed reduction in spontaneous GFP activation in GSCs and efficient GFP activation in GB/SGs suggests that the Bam-FLPD5 in combination with nos>>mCD8GFP detects differentiating cells with higher specificity.

### Quantifying specificity and efficiency of lineage marking

To estimate the number of GFP-positive cells that reflect true dedifferentiation events, we formulated a computational model using observed rates of key cellular processes in our previous live imaging study, including: asymmetric division (*p*), symmetric self-renewal (*q*), symmetric differentiation (*r*), dedifferentiation (*k*_*dd*_). Then, we defined flippase recombination rate in GB/SGs to be (⍺) and recombination occurring in GSCs (⍺*) as the spontaneous background event, disregarding stage-specific promoter activity. Because this background recombination must also occur at the same rate in GB/SGs, the stage-specific recombination rate in GB/SGs (⍺^True^) is determined as (⍺^True^ = ⍺ - ⍺*) (all rates are per cell cycle, Figure S1A, B).

Using a publicly available software *Virtual Cell (*VCell) [25, 26], we first simulated the time-dependent changes in the fraction of GFP-positive GSCs and validated the model by fitting the curve to the experimentally observed GFP rates. For the data obtained by using both nos>>GFP and nos>>mCD8GFP cassettes, the model outputs closely matched the experimental data (Figure 4A, B), yielding values of ⍺* that are consistent with the estimates obtained directly from the experimental data, see methods section “Estimation of flippase recombination rate in GSCs (⍺*)” for details. These validate the model and indicate its ability to accurately recapitulate the frequency of marked cells in the niche.

**Figure 4.**
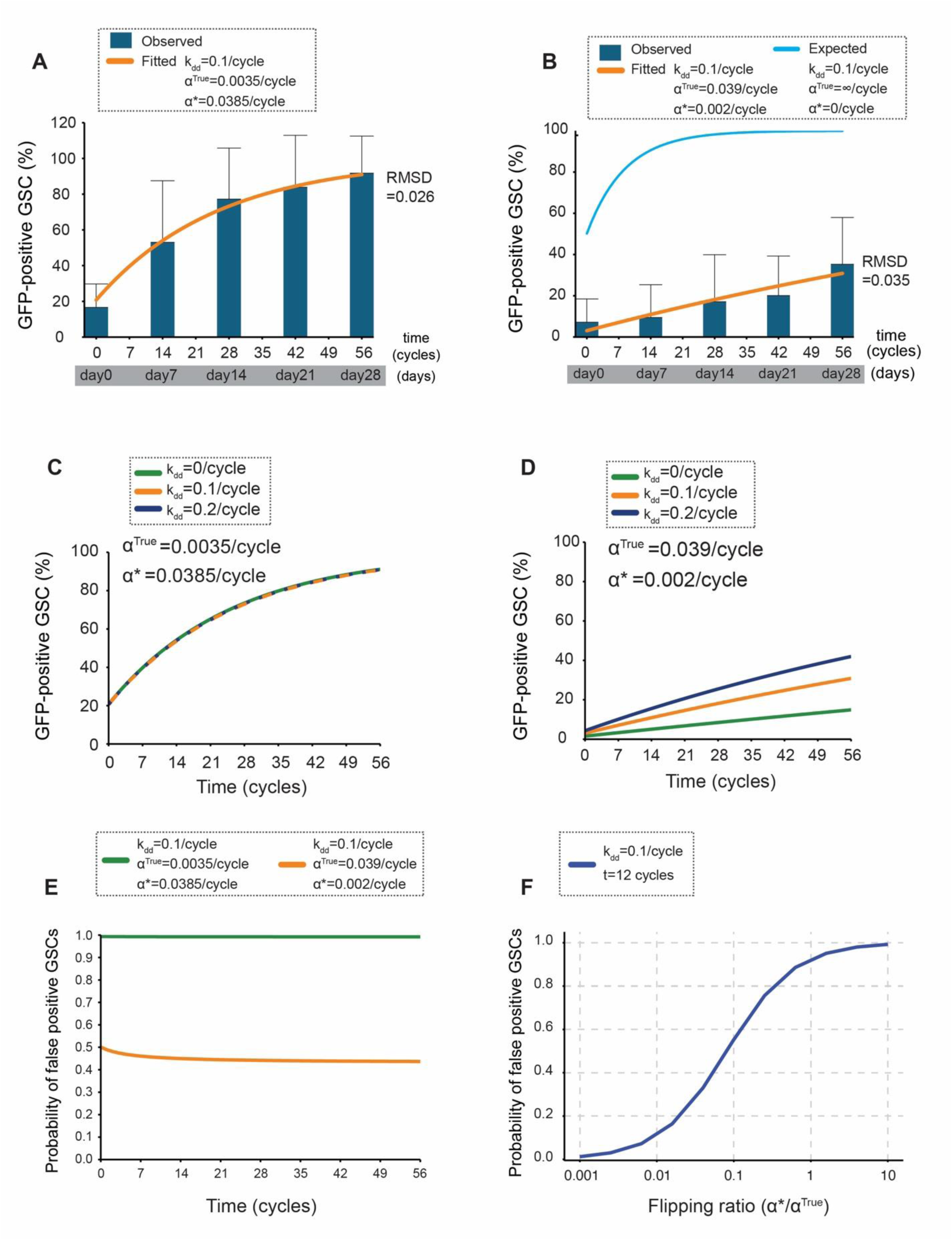
Quantifying specificity and efficiency of lineage marking. **A)**Model fitting for the Bam-FLPD5, nos>>GFP system. Experimental data showing the percentage of GFP-positive GSCs over time (blue bars; data from Figure 2D) are compared to the best-fit model output (orange curve). The model incorporates key cellular processes (see Figure S1A, B and Materials and Methods for details). The root mean squared deviation (RMSD) between the model and the experimental data is 0.026. Two cycles are equivalent to one day [46]. Fitted parameters (⍺^True^ and ⍺*) are shown in the graph. **B)** Model fitting for the Bam-FLPD5, nos>>mCD8GFP system. Experimental data showing the percentage of GFP-positive GSCs over time (blue bars; data from Fig 3C) are compared to the best-fit model output (orange curve). The root mean squared deviation (RMSD) between the model and the experimental data is 0.035. The blue line represents the expected percentage of GFP-positive GSCs if the recombination efficiency was 100% in all differentiating cells (⍺^True^=∞) and with no spontaneous GFP activation (⍺*=0) **C)** Predicted effect of dedifferentiation rate (*k*_*dd*_) on GFP-positive GSCs in the Bam-FLPD5, nos>>GFP system. Model predictions for the percentage of GFP-positive GSCs over time are shown for different dedifferentiation rates (*k*_*dd*_ = 0, 0.1, and 0.2 per cell cycle). The overlapping lines demonstrate that the system is insensitive to changes in *k*_*dd*_ due to high ratio of ⍺*/⍺^True^ = 11. **D)** Predicted effect of dedifferentiation rate (*k*_*dd*_) on GFP-positive GSCs in the Bam-FLPD5, nos>>mCD8GFP system. Model predictions for the percentage of GFP-positive GSCs over time are shown for different dedifferentiation rates (*k*_*dd*_ = 0, 0.1, and 0.2 per cell cycle). The lines demonstrate the system’s sensitivity to changes in *k*_*dd*_ due to lower ratio of ⍺*/⍺^True^ = 0.0667. **E)** Predicted probability of false positive GSCs over time. The model-predicted probabilities of a GFP-positive GSC being a false positive are shown for the Bam-FLPD5, nos>>GFP system (green line) and the Bam-FLPD5, nos>>mCD8GFP system (orange line). The nos>>GFP system shows a near 100% false positive probability, while the nos>>mCD8GFP system shows a significantly lower probability of approximately 45%. **F)** Predicted probability of false positive GSCs as a function of the flipping ratio over time for two different marking systems with varying flipping ratios (⍺*/⍺^True^). The analysis was performed at a representative time point (t=12 cycles).

### Stage specificity of the recombination determines the feasibility of the detection system

We next used the model to estimated how many of GFP positive cells are true dedifferentiation cells. When the nos>>GFP cassette was used, spontaneous flippase recombination in GSCs occurs at a rate nearly equal to the flippase recombination in GB/SGs, ⍺* ≈ ⍺, indicating that the flippase activity is entirely not stage specific. In this case, the model revealed that dedifferentiation rate *k*_*dd*_ had no discernible effect on the frequency of GFP-positive GSCs, as expected. Indeed, the predicted frequency of GFP-positive GSCs remained essentially same across a range of *k*_*dd*_ values (0, 0.1, and 0.2) (Figure 4C), indicating that the observed GFP-positive rate is not reporting the dedifferentiation rate at all.

In contrast, when nos>>mCD8GFP cassette was used, which has a significantly lower flippase recombination rate in GSCs than in GB/SGs, ⍺*<< α, the GFP positive rate showed the clear sensitivity to the dedifferentiation rate *k*_*dd*_, as the percentage of GFP-positive GSCs significantly increases with the increasing *k*_*dd*_ (Figure 4D). These results highlight the importance of the stage specificity of flippase excision events in marking the cells for estimating dedifferentiation frequency.

An important aspect of lineage tracing is distinguishing true lineage-marked cells from those marked due to non-specific flipping events (false positives). The model estimates the rate of ‘false positives’ as GFP-positive GSCs originating from two sources: 1) GSCs that spontaneously activate GFP within the niche and 2) dedifferentiated GFP-positive GSCs whose lineage traces back to a spontaneous GFP activation event within and outside the niche. With the Bam-FLPD5, nos>>GFP system, the model revealed that essentially all (∼100%) of the observed GFP-positive GSCs at all time points were false positives (Figure 4E, green line). In contrast, with the nos>>mCD8GFP cassette, the fraction of false positives was reduced to about 45% across all time points (Figure 4E, orange line). While still substantial, this represents a significant improvement over the Bam-FLPD5, nos>>GFP system.

To further explore the relationship between the stage specificity of the flipping event and false positive rates, we systematically varied the “flipping” ratio (⍺*/⍺^True^) – the ratio of background flippase activity in GSCs (⍺*) to the stage-specific activity in GB/SGs (⍺^True^) – while keeping other parameters constant. We performed this analysis at a representative time point (t=12 cycles), given that the probability of false positives is largely time-independent (Figure 4E). As expected, low flipping ratios yield low false-positive rates approaching 0, but as the flipping ratio increases, the probability of false grows rapidly, showing a sigmoid semi-log dependence (Figure 4F). These findings indicate that minimizing background flippase activity is a critical factor for the genetic marking system to be successful.

In summary, our modeling analysis demonstrates that the Bam-FLPD5, nos>>mCD8GFP system reports dedifferentiation rates much more efficiently than the previous system, although approximately half of the GFP positive GSCs are still false positives. The expected “true” dedifferentiation rate, assuming zero background recombination in GSCs, and 100% recombination in GB/SGs, is represented by the blue line in Figure 4B. This curve indicates that the dedifferentiated cells rapidly occupy the entire niche within ∼14 days of fly’s age (Figure 4B). The difference between the blue and orange curves represents the false negative rate that needs to be improved in future studies.

### Experimentally increasing the dedifferentiation rates results in a high frequency of marked GSCs

To experimentally test the model’s predicted sensitivities to dedifferentiation, we introduced a heat-shock-inducible Bam (hs-Bam) transgene into flies carrying either the Bam-FLPD5, nos>>GFP or Bam-FLPD5, nos>>mCD8GFP lineage marking system. The hs-Bam transgene is commonly used to induce dedifferentiation; a heat-shock treatment temporarily forces GSCs to differentiate, and when treatment stops, GSCs are replenished through dedifferentiation [27].

Our model predicts that the Bam-FLPD5, nos>>GFP system, with its high ⍺*/⍺^True^ ratio, would be insensitive to changes in *k*_*dd*_ (Figure 4C), while the Bam-FLPD5, nos>>mCD8GFP system, with its lower ⍺*/⍺^True^ ratio, would show an increase in GFP-positive GSCs with increasing *k*_*dd*_ (Figure 4D).

At 7 days after eclosion, during which the flies were kept at room temperature, we observed a significant increase in GFP-positive GSCs in flies carrying the hs-Bam transgene with the Bam-FLPD5, nos>>mCD8GFP system (Figure 5A, B, E). This is likely due to the basal activity of the heat-shock promoter at room temperature, leading to continuous Bam expression. This continuous Bam expression causes GSC loss, which subsequently induces the replacement events likely through dedifferentiation. Notably, this increase in GFP-positive GSCs is consistent with the model’s prediction that Bam-FLPD5, nos>>mCD8GFP system is sensitive to changes in the dedifferentiation rate.

**Figure 5.**
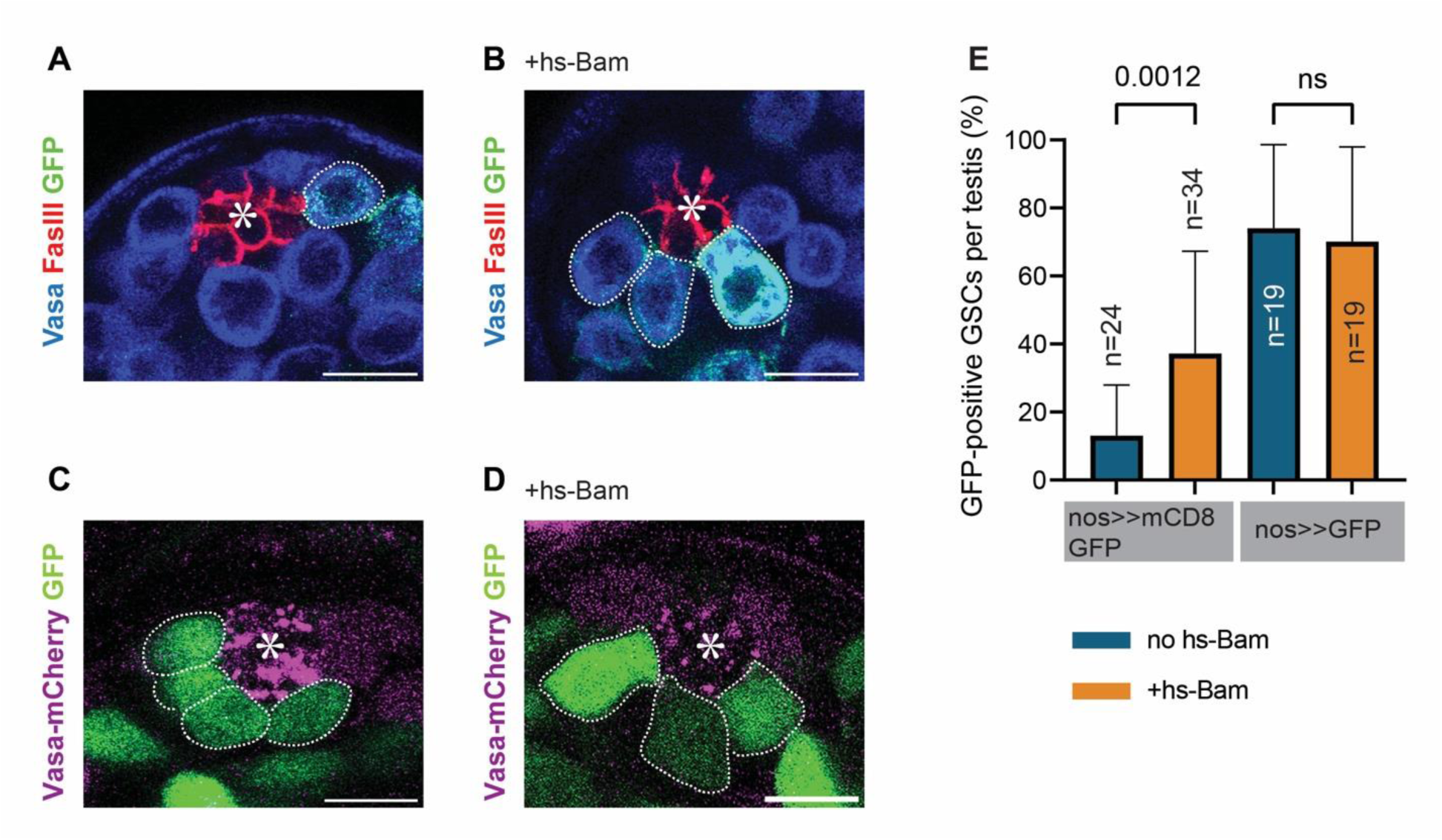
Introduction of a hs-Bam transgene results in high frequency of marked GSCs. **A, B)** Representative immunofluorescence images of the testis tip from 7-day-old flies, kept at room temperature, with Bam-FLPD5 marking system with the nos>>mCD8GFP cassette alone **(A)** or with an additional single copy of the hs-Bam transgene **(B)**. Testes were stained with Vasa (blue) and FasIII (red) antibodies. GFP-positive GSCs (green) are outlined with white dotted lines. **C, D)** Representative images of live testis tip from 7-day-old flies, kept at room temperature, with Bam-FLPD5 marking system with the nos>>GFP cassette alone **(C)** or a combination with a single copy of hs-Bam **(D)**. **E)** The graph shows the quantification of GFP-positive GSCs per testis in 7-day-old flies, kept at room temperature, for the indicated genotypes. All scale bars are 10 μm. Asterisks in the images indicate the approximate location of the hub. p-values were calculated by Šídák’s multiple comparisons test. “n” indicates the number of testes scored.

In contrast, and as predicted by our model, increasing dedifferentiation frequency in flies carrying the hs-Bam transgene with Bam-FLPD5, nos>>GFP system, did not result in a significant change in GFP-positive GSCs under the same conditions (Figure 5C-E). These data suggest that the observed increase in GFP-positive GSCs in the Bam-FLPD5, nos>>mCD8GFP system reflects the dedifferentiation events.

Interestingly, the standard heat-shock protocol to induce dedifferentiation with hs-Bam at 37°C followed by a 3 to 7-day recovery period, did not result in a significant change in GFP-positive GSCs with both nos>>GFP and nos>>mCD8GFP cassettes (Figure S3). We speculate that high levels of Bam production during heat-shock treatments may cause negative feedback mechanisms that suppress transcription from the Bam promoter used for the FLPD5 (Bam-FLPD5), thereby reducing the efficiency of marking differentiating cells.

## Discussion

Lineage tracing is a powerful tool for tracking cellular dynamics *in vivo*, including proliferation, differentiation, and dedifferentiation. Recombinase-based systems, such as Cre-loxP and FLP-FRT are widely used in lineage tracing. However, their reliance on tissue-specific promoters makes them susceptible to “leaky” or off-target recombination. Studies in mice have demonstrated the non-specificity of various Cre lines [28, 29]. Ideally, a lineage-specific promoter should be distinctly active (“ON”) in one target lineage and inactive (“OFF”) in all other lineages.

In this study, we developed an enhanced genetic marking system (Bam-FLPD5) to identify dedifferentiated GSCs with improved sensitivity. By addressing key limitations of previous lineage tracing methods, including delayed reporter activation that reduces marking efficiency, our Bam-FLPD5 system offers a more accurate approach for monitoring dedifferentiation events. While genetic marking is a valuable tool for studying dedifferentiation, live imaging remains essential for confirming these events and distinguishing them from artifacts like spontaneous reporter activation in off-target cells. Several past studies have utilized live imaging to observe stem cell dynamics in real time, including studies in mouse spermatogonial stem cells [30], and *Drosophila* male GSCs [31]. Through real-time tracing of cells within intact tissues, these studies have provided insights into cellular plasticity and ability of partially or terminally differentiated cells to revert to a stem cell state. In this study, we integrated live imaging with our genetic marking approach to further enhance the accuracy and provide an additional layer of validation.

Our findings emphasize spontaneous recombination’s significant and previously underestimated impact on lineage tracing studies. Spontaneous recombination likely occurs in other lineage tracing systems but has often been overlooked due to the lack of real-time validation methods like live imaging. Our simulations showed a strong positive correlation between the false positive rates and the FLP recombination ratio (⍺*/⍺) (Figure 4F). For example, the Bam-FLPD5, nos>>GFP system, where the estimated ⍺*/⍺ ratio was ∼1 (almost no stage specificity of the recombination), our simulation showed essentially all of GFP-positive GSCs (∼100%) across 56 cycles are false positives as we expected. This suggests that none of the observed GFP-positive GSCs in this system represent the true dedifferentiation events (Figure 4E, green line). In contrast, the Bam-FLPD5, nos>>mCD8GFP system, with a lower ⍺*/⍺ ratio, exhibited a reduced false positive rate of approximately 45% (Figure 4E, orange line), implying that roughly 55% of GFP-positive GSCs likely represent true dedifferentiation. This enables the use of this system to monitor the dedifferentiation rate for future studies.

Although our new genetic marking system improved the sensitivity of previously established systems, it still does not capture all dedifferentiated cells. A comparison between the observed percentages of dedifferentiated GSCs in the Bam-FLPD5 system (Figure 4B, blue bars) and the expected curve derived from our mathematical model (Figure 4B, blue line), which assumes 100% recombination efficiency in all differentiating cells, reveals that a large portion of differentiating cells remains unmarked with currently achieved recombination efficiency. There is still a need for further research focused on identifying gene promoters with both high activity and high specificity.

With the advances in next-generation sequencing and single-cell technologies, computational-based lineage tracing tools have emerged (reviewed in [32, 33]). These include Polylox DNA barcoding, which uses Cre recombinase to create diverse combinations barcodes to uniquely label cells of interest for lineage tracing (although promoter specificity of Cre expression remains a consideration) [34, 35]. Other approaches leverage naturally occurring somatic mutations, such as those in mitochondrial DNA (mtDNA) [36], or engineered mutations introduced via CRISPR-Cas9 systems [37] to infer cellular relationships. While DNA barcoding and marker-independent methods offer valuable insights, they often lack the ability to directly validate the lineage tracing events in real-time. A key advantage of our approach is the integration of long-term live imaging with genetic marking. This combination allows for direct assessment of accuracy, a capability lacking in many newer techniques.

GSCs typically divide asymmetrically, with several intrinsic factors non-randomly segregated between the stem cell and its differentiating daughter cell [38–40]. However, dedifferentiation disrupts this intrinsic asymmetry, and whether this results in differences between original and dedifferentiated stem cells is not well understood. Moreover, dedifferentiated stem cells have been implicated as a cellular origin of tumorigenesis in several tissues [41, 42]. Lineage tracing tools, such as the Bam-FLPD5 system developed in this study, provide a valuable approach to investigate these questions.

## Materials and methods

### Fly husbandry and strains

Flies were raised on standard Bloomington medium at 25°C (unless temperature control was required). 0- to 7-day-old adults were used for all experiments unless specific ages of flies were noted. The following fly stocks were obtained from Bloomington Stock Center (BDSC); nosGal4 (BDSC64277) [43]; 20XUAS-FLPD5 (BDSC55805); UAS(FRT.stop)mCD8-GFP (BDSC30032); hs-flp (BDSC1929) and hs-bam (BDSC24636). pVas-Vasa mCherry (FBtp0065762) [44] was a kind gift from Yukiko Yamashita. nos-FRT-mCherry-stop-FRT-Gal4, UAS-GFP [23] was from Inaba lab stock.

### Immunofluorescence staining

Testes were dissected in 1x phosphate-buffered saline (PBS) (Fisher Scientific) and fixed in 4% formaldehyde (Electron Microscopy Sciences) in PBS for 30 minutes. Following fixation, testes were washed in PBS containing 0.2% TritonX-100 (Fisher Scientific) (PBST) for a minimum of 60 minutes (three washes of 20 minutes each). They were then incubated overnight at 4°C with primary antibody in 3% bovine serum albumin (BSA) (Sigma) in PBST. The following day, samples were washed in PBST for 60 minutes (three washes of 20 minutes each), incubated with secondary antibody in 3% BSA in PBST at room temperature for 2 hours, and then washed again in PBST for 60 minutes (three washes of 20 minutes each). Finally, samples were mounted using VECTASHIELD with 4’,6-diamidino-2-phenylindole (DAPI) (Vector Lab).

The primary antibodies used were as follows: rat anti-Vasa (1:20; developed by A. Spradling and D. Williams, obtained from Developmental Studies Hybridoma Bank (DSHB); mouse anti-hu-li tai shao (Hts) (1:20, 1B1; DSHB) and mouse-anti-Fasciculin III (FasIII) (1:40, 7G10; DSHB). AlexaFluor-conjugated secondary antibodies (Abcam) were used at a dilution of 1:400.

### Long-term live imaging

Long-term live imaging was conducted using a previously established method [31]. Testes were dissected in 1X Becker Ringer’s [31] solution and mounted onto 35mm Glass Bottom Dishes (Nunc). To prepare the imaging dish, 500 μL of 1 mg/mL poly-L-lysine (Sigma) was applied to the coverslip area of the dish and incubated at room temperature for 5- to 7-hours. Following incubation, the poly-L-lysine solution was replaced with the Becker Ringer’s solution. Testes were then dissected in Becker Ringer’s solution and mounted on the poly-L-lysine coated surface with the tips oriented downward. Next, Becker Ringer’s solution was carefully removed, and pre-warmed (room temperature) Schneider’s *Drosophila* medium (supplemented with 10% fetal bovine serum and glutamine–penicillin–streptomycin, Sigma) was added to the dish.

Z-stack images were acquired at 2 µm intervals (10-15 stacks total) using the Zeiss LSM800 Airyscan microscope with a 63× oil immersion objective (NA = 1.4) every 10 minutes over a 16-hour period. The preset tiling function was utilized for automated image acquisition.

### Image analyses

Airyscan time-lapse images were processed with Zen software (Zeiss). Each z-stack was manually analyzed to track cells, record their GFP-expression status, and identify dedifferentiation events. The hub area was identified as a circular, autofluorescent region located near the testis tip, surrounded by fluorescently labeled germline stem cells. Dedifferentiation events were recorded when germline cells, initially positioned more than one cell layer away from the hub’s edge reattached to the hub and maintained their attachment for at least 30 minutes. The moment a dedifferentiated cell observed at the hub edge was marked as the “reattachment” time point. Movies were annotated using Veed.io.

### Estimation of flippase recombination rate in GSCs (⍺*)

The flippase recombination rate in GSCs (⍺*) was determined directly from live-imaging data. For the BamFLPD5, nos>>GFP system, 27 testes were imaged for 12 hours each (Figure 2E). Each testis contains an average of 9 GSCs. Across all testes, 11 GSCs were observed to become GFP-positive during the imaging period. Therefore, ⍺* was calculated by dividing the total number of flipping events in GSCs (11) by the total number of GSC cell cycles observed (27 testes*9 GSCs/testis=243 GSC cycles), yielding a value of 0.045 flipping events per cell cycle. A similar procedure was followed for the Bam-FLPD5, nos>>mCD8GFP system, using its respective experimental data in Figure 3D. Here, ⍺* was calculated by dividing the total number of flipping events in GSCs (1) by the total number of GSC cell cycles observed (25 testes*9 GSCs/testis=225 GSC cycles), yielding a value of 0.004 flipping events per cell cycle.

### Short-term live imaging

Short-term live imaging was performed for static image acquisition of testes to avoid potential signal loss or loss of extracellularly located fluorescent protein (GFP) during fixation and permeabilization. Dissected testes were placed in Schneider’s Drosophila medium (supplemented with 10% fetal bovine serum and glutamine–penicillin–streptomycin, Sigma), then placed onto two etched rings on Gold Seal Rite-On Micro Slides with media and covered with coverslips. Z-stack images were acquired at 1 µm intervals (15-20 stacks total) using the Zeiss LSM800 Airyscan microscope with a 63× oil immersion objective (NA = 1.4). For these short-term live imaging experiments, all images were taken within 30 minutes of dissection, without time-lapse imaging.

### GSC depletion and recovery

GSC depletion and recovery were performed as previously described, with modifications [27]. Adult flies (0- to 3-days old) carrying the hs-Bam transgene (BDSC24636) were raised at 22°C. They underwent six heat shock treatments in a 37°C water bath for 30 minutes, twice daily, within vials with fly food. Between treatments, the vials were maintained in a 29°C incubator. Following the last heat shock treatment, the flies were returned to 22°C for the recovery phase. Testes were dissected after a 5-day recovery period.

### Generation of Bam-FLPD5 transgenic line

The minimum bam promoter sequence (-198 to transcription start site, TSS) [45] was amplified from genomic DNA isolated from wild-type flies using the following primers:

SphI KpnI bam -198 Forward: 5’- AGCGGATCCAAGCTTGCATGCGGTACCCCAAATCAGTGTGTATAATT-3’ Bam -198 Reverse: 5’-TATTCTTAAGTTAAATCACACAAATCACTCGAT-3’

Separately, the FLPD5 sequence, excluding the PEST sequence, was amplified from the genomic DNA of the 20XUAS-FLPD5 strain (BDSC55805) using the following primers:

FLPD5 Forward: 5’- GTGATTTGTGTGATTTAACTTAAGAATAATGCCGCAGTTTGATATCCTCTG-3’ FLPD5 Reverse: 5’-GTACCCTCGAGCCGCTTAAATACGGCGATTGATGTAGGAGCTCA- 3’

pUAST-attB vector was digested by SphI and NotI restriction enzymes to remove the UASt promoter. The amplified bam -198 and FLPD5 fragments were assembled with the digested vector using a Gibson Assembly kit (NEB) and sequenced. Transgenic flies were generated using strain attP2 by PhiC31 integrase-mediated transgenesis (BestGene).

### Mathematical modeling

#### Model

The model was developed for the analysis of efficiency, specificity, and sensitivity of genetic constructs designed for the identification of dedifferentiated germline stem cells (GSCs) in *Drosophila* testes.

The model outputs must mimic experimental readouts - time-dependent fractions of marked dedifferentiated GSCs. Correspondingly, we formulate our model in terms of numbers of unmarked and marked GSCs in the niche and corresponding GBs and SGs available for dedifferentiation in the vicinity of the niche. The stem cell pool is presented in the model by the following variables: the number of the innate unmarked stem cells, *n*_0_(*t*); the number of the dedifferentiated unmarked stem cells, *n*_1_(*t*); the number of the dedifferentiated stem cells marked as a result of the promoter activation and subsequent flippase recombination, *n*_2_(*t*); and the number of the stem cells marked by spontaneous nonspecific GFP activation (false positives), *n*_3_(*t*). The GB/SGs pool is described by the following variables: the number of unmarked GB/SGs, *m*_1_(*t*); the number of GB/SGs marked due to the promoter activation, *m*_2_(*t*); and the number of GB/SGs marked by spontaneous GFP activation, *m*_3_(*t*).

The variables described above are governed by the following key processes included in the model (Figure S1A,B): asymmetric division (characterized by rate constant *p*), symmetric renewal (*q*), symmetric differentiation/loss (*r*), dedifferentiation of GB/SGs (*k*_*dd*_), differentiation of GB/SGs (β), flippase recombination in GB/SGs (⍺), flippase recombination in GSCs (⍺*), and stage-specific flippase recombination in GB/SGs (⍺^True^) (⍺^True^ = ⍺ - ⍺*) (all rates are per cell cycle).

We assume that the processes involving unmarked and marked cells are described by equivalent parameter values. We further assume that the niche tightly controls the total number of GSCs, *N* = *n*_0_(*t*) + *n*_1_(*t*) + *n*_2_(*t*) + *n*_3_(*t*), so that it remains constant at all times,

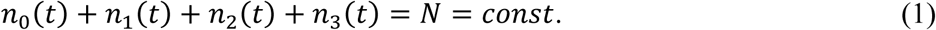

Experimental fractions of GFP-positive cells were obtained from the pools containing large numbers of cells for each time point (on the order of a hundred or more), so that the fluctuations of their averages were relatively small. This allowed us to ignore the discreteness of the variables and approximate the dynamics of the corresponding fractions, 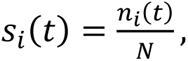 *i* = 0,1,2,3, and 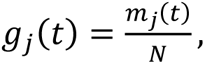 *j* = 1,2,3, by differential equations:

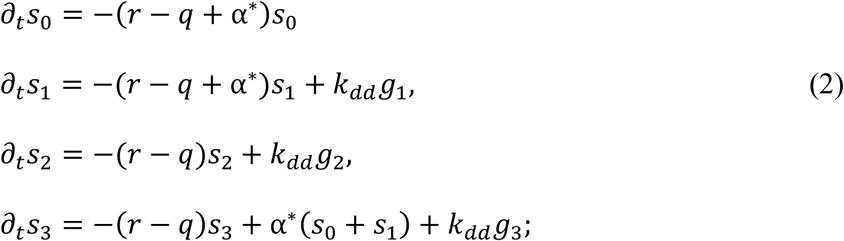

and

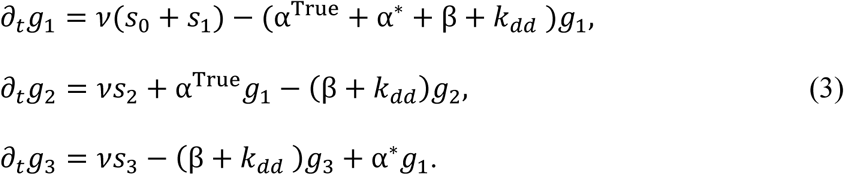

Parameter *v* in Eqs (3) represents the stem-cell mitotic rate, *v* = 1 per cell cycle.

The system of Eqs (2) and (3) was solved using VCell (www.vcell.org). The VCell implementation of the model can be found in the VCell database of public MathModels under username “boris”; the model name is “Inaba_dedifferentiation_model_public”.

#### Model Parameters

We constrained model parameters using the results of long-term live imaging, along with the analysis of the model’s salient features.

Not all parameters of the model are independent. Experimental observations indicate that by the time of eclosion, the pool of differentiated cells available for dedifferentiation has approached its steady-state size equaling *N*, i.e. *m*_1_(*t*) + *m*_2_(*t*) + *m*_3_(*t*) = *N* = *const* and 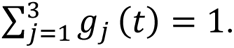 This makes the mechanism of replenishing of GSCs by dedifferentiation mathematically equivalent to symmetric renewal with an effective *q*_*eff*_ = *k*_*dd*_ + *q* = *r* and *p* + *q*_*eff*_ + *r* = *v*. Indeed, from Eq (1), 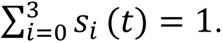 Then summing up Eqs (2) yields 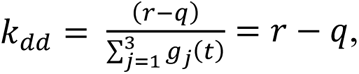 and since 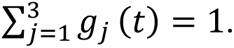

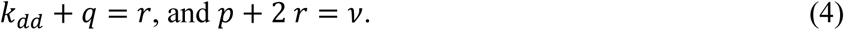

Eqs (4) are consistent with the data cited at the beginning of the section *Improvement of genetic marking system for identifying dedifferentiated GSCs*. Correspondingly, *p* = 0.72/cycle and *k*_*dd*_ = 0.1/cycle, then 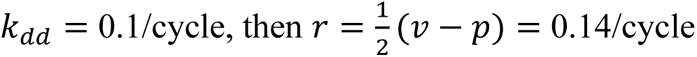 and *q* = *r* − *k*_*dd*_ = 0.04/cycle.

Similarly, summing up Eqs (3) yields *v* = β + *k*_*dd*_, and β = 0.9/cycle.

#### Fitting the model to experimental data

To determine parameters ⍺^True^ and ⍺*, we fitted our model to experimental time dependences of fractions of the GFP-positive GSCs obtained for a given genetic marker with an experimentally constrained ratio ⍺*/⍺^True^. For this, Eqs (2) and (3) should be solved with varying ⍺^True^ and initial conditions, to minimize the deviation between the solution of the model and the experimental data. As the experimental data are available beginning with the fly’s eclosion, trying possible values of the seven model variables at the time of eclosion would be unwieldy. Instead, we moved the zero time point to the time in the larvae stage at which stem cells start to divide. It is reasonable to assume that at that time, all GCSs are unmarked, and the numbers of differentiated cells are zero. This allowed us to solve the model with same initial conditions in all trials and only vary the time of eclosion, *τ*_*ecl*_. One caveat is that 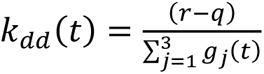 becomes now a function of time, since 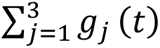 is not at steady state at *t* ≤ *τ*_*ecl*_. To avoid division by zero in the equation for *k*_*dd*_(*t*), the initial conditions used in all our trials were as follows: *s*_0_(0) = 1, *s*_*i*_(0) = 0 (*i* = 1,2,3), *g*_0_(0) = ε, *g*_*j*_(0) = 0 (*j* = 2,3), where ε is a small number (in our simulations, *ε* = 10^-6^). The results were then offset by *τ*_*ecl*_ for comparison with the experimental data.

We characterized discrepancy between the simulation results and experimental data by the root-mean-square deviation (RMSD) defined as 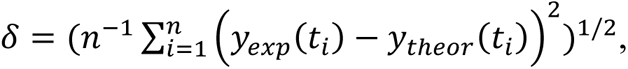 where *n* is the number of time points, for which we performed the comparison, *y*_*exp*_(*t*_*i*_) are the experimental values at times *t*_*i*_, and *y*_*theor*_(*t*_*i*_) = *s*_2_(*t*_*i*_) + *s*_3_(*t*_*i*_) ≡ *s*_+_(*t*_*i*_). Good fits, characterized by *δ*∼ 0.02-0.04, were obtained with *τ*_*ecl*_ = 6 cycles.

Once fully parameterized, the model was used to explore sensitivities of the marking systems to *k*_dd_ and ⍺^true^, and to compute the probability of false positives, 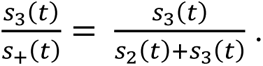

### Statistical analysis and graphing

No statistical methods were used to predetermine the sample size. The experiment values were not randomized. The investigators were not blinded to allocation during experiments and outcome assessment. All experiments were independently repeated at least 3 times to confirm the results. Statistical analysis and graphing were performed using GraphPad prism9. The illustrations were generated by using Adobe Illustrator or BioRender. All data are shown as means ± s.d.

## Data Availability

The data that support all experimental findings of this study are available within the paper and its Supplementary Information files and in the BioStudies database under the accession number S-BSST1903 (https://www.ebi.ac.uk/biostudies/studies/S-BSST1903). Source_data and Simulation_source_data contain numerical values for each experiment and simulations.

## Acknowledgments

We thank Yukiko M. Yamashita, Michael Buszczak and the Bloomington *Drosophila* Stock Center, and the Developmental Studies Hybridoma Bank for reagents. This research is supported by R35GM128678 from the National Institute for General Medical Sciences and start-up funds from UConn Health (to M.I.). B.M.S. was supported in part by R24GM137787 from National Institute for General Medical Sciences.

## Author Contributions

M.I. and M.B.B. conceived the project, designed and executed experiments and analyzed data. B.M.S. and M.B.B. conducted mathematical modeling and data fitting using *Virtual Cell*. All authors wrote and edited the manuscript.

