## Supplemental Figure for "Detection of dedifferentiated stem cells in *Drosophila* testis"

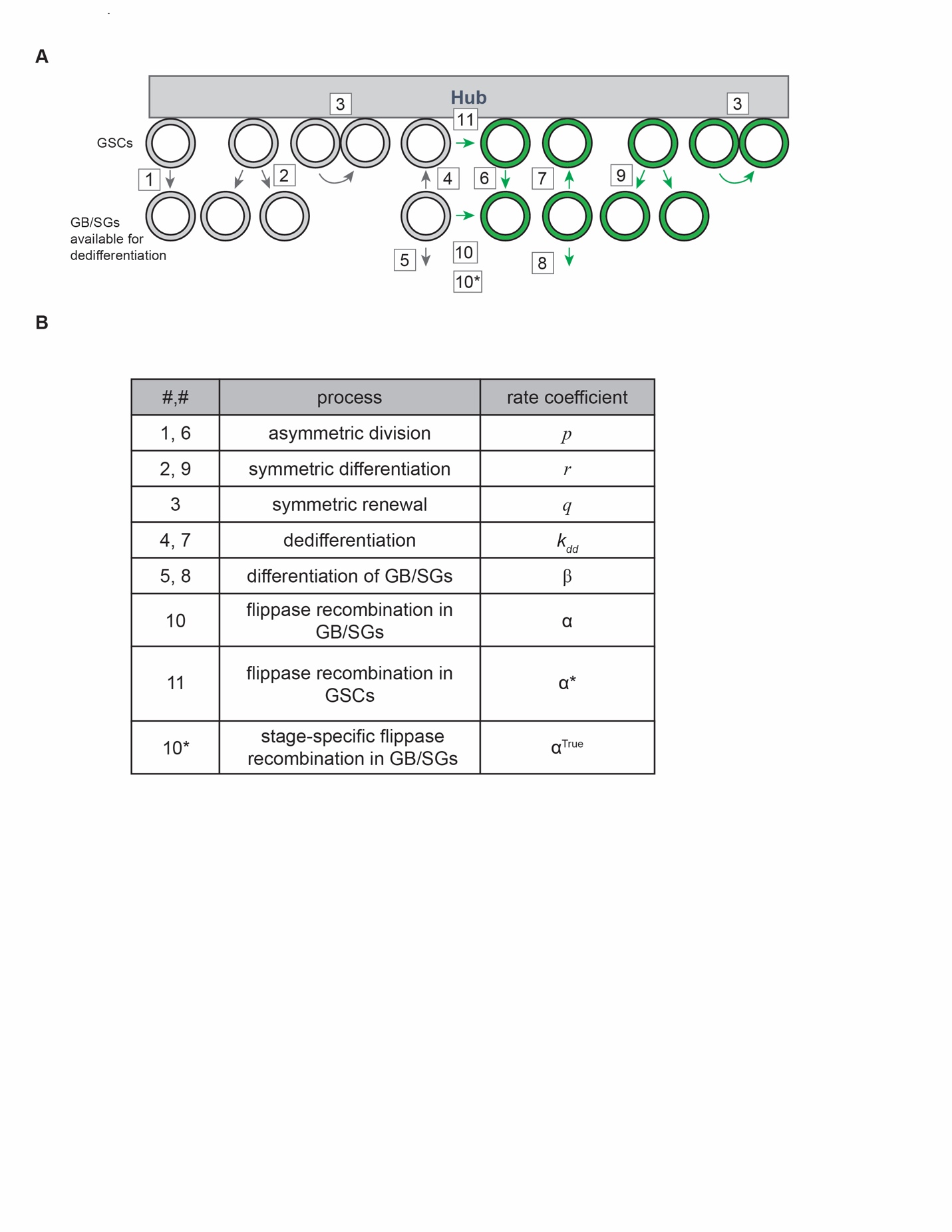

**Figure S1. Processes and rate coefficients used in the mathematical modeling.**

**A)** Schematic diagram of GSC niche dynamics, illustrating the cellular processes included in the mathematical model. GSCs (top row) are adjacent to the hub. GBs and SGs (bottom row) are differentiating cells. Arrows represent cellular processes, with numbers corresponding to the processes listed in panel B.

**B)** The table lists the cellular processes considered in the model, along with the symbol used to represent their respective rate coefficients. These processes include: asymmetric division (*p*), symmetric renewal (*q*), symmetric differentiation/loss ($r$), dedifferentiation of GB/SGs ($k_{dd}$), differentiation of GBs/SGs (β), flippase recombination in GB/SGs (⍺), flippase recombination in GSCs (⍺*), and stage-specific flippase recombination in GB/SGs (⍺^True^). Further details on the mathematical model and its analysis are provided in the ‘Mathematical modeling’ subsection of the *Materials and Methods*).

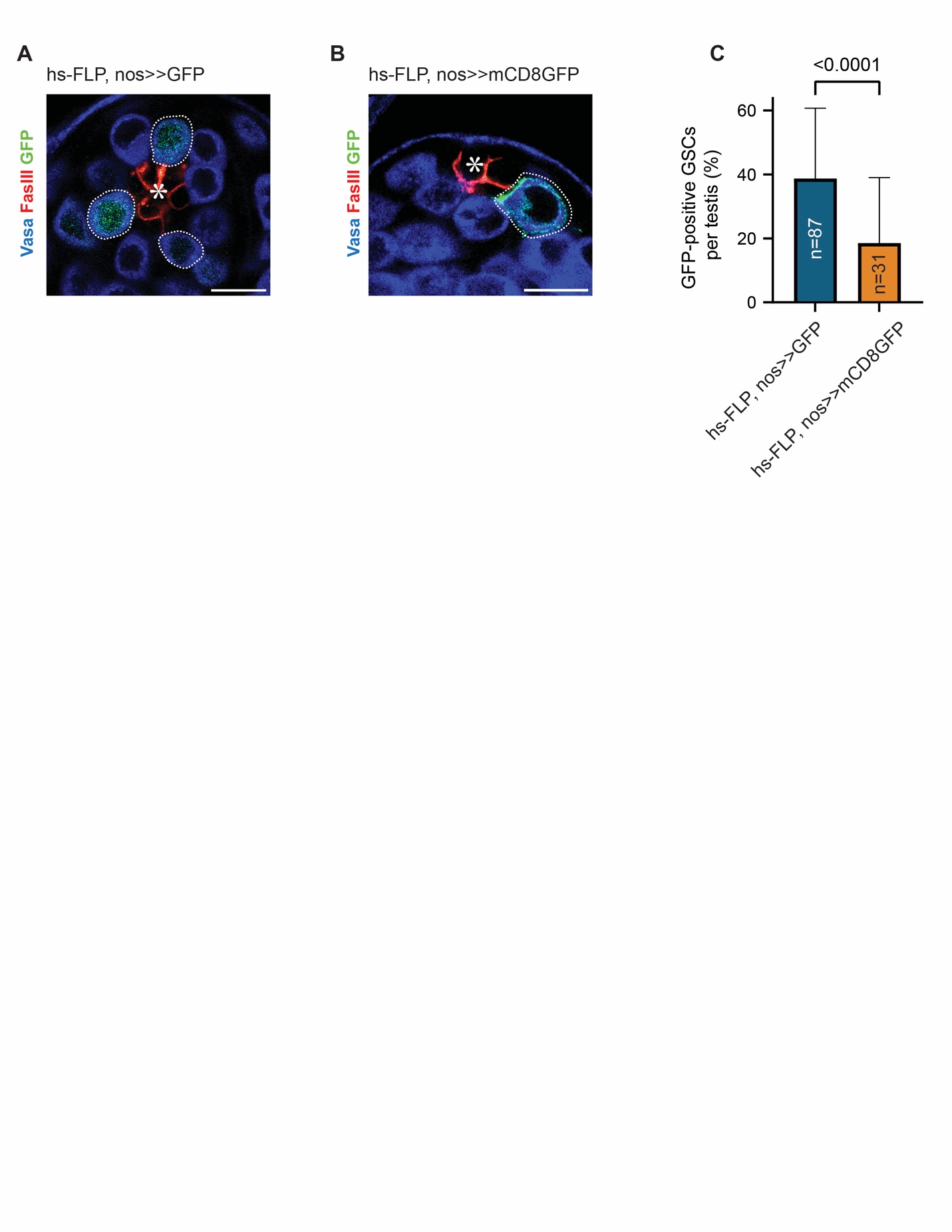

**Figure S2. The nos>>mCD8GFP cassette has a higher threshold for FLP-mediated recombination than the nos>>GFP cassette in GSCs.**

**A, B)** Representative immunofluorescence images of the testicular niche 24 hours after a single 30-minute of heat-shock in flies with hs-FLP, nos>>GFP genotype **(A)** and the hs-FLP, nos>>mCD8GFP genotype **(B)**. Testes were stained with Vasa (blue) and FasIII (red). GFP-positive GSCs (green) are encircled by white dotted lines.

**C)** The graph shows the percentage of GFP-positive GSCs per testis 24 hours after a single 30-minute of heat-shock for the indicated genotypes.

All scale bars are 10 μm. The hub is marked by asterisks. The p-value was calculated by two-tailed student t-test. “n” indicates the number of testes scored.

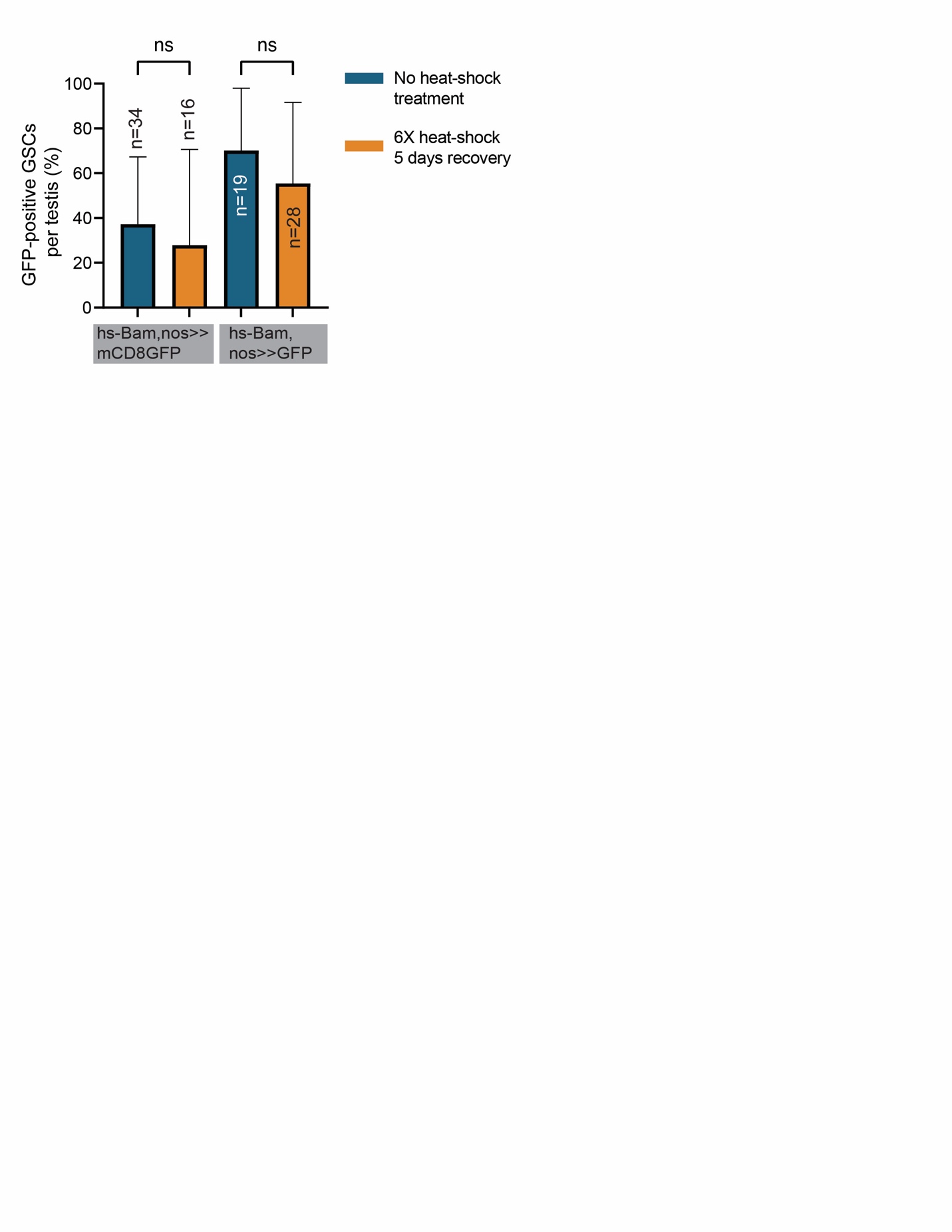

**Figure S3. Heat-shock induced Bam overexpression reduces recombination efficiency of Bam promoter driven flippase.**

Quantification of GFP-positive GSCs per testis in flies subjected to either no heat-shock treatment (blue bars) or six heat-shock treatments at 37ºC followed by a 5-day recovery period (orange bars) using both nos>>mCD8GFP and nos>>GFP cassettes combined with Bam-FLPD5 and a single copy of the hs-Bam transgene. The no heat-shock treatment data (blue bars, hs-Bam) are the same data presented in Figure 5E and represent flies kept at room temperature for 7 days post-eclosion to match the total experimental duration of the heat-shock treatment group. P-values were calculated by Šídák's multiple comparisons test. “n” indicates the number of testes scored.

**Movie legends**

**Movie S1: Spontaneous GFP activation in a GSC expressing Bam-FLPD5, nos>>GFP marking system**

Time-lapse movie of a testis tip showing the spontaneous GFP-activation in a GSC (corresponding to Figure 2E). The testes are expressing Bam-FLPD5, nos>>mCD8GFP marking system, and germline cells are visualized by Vasa-mCherry (magenta). The arrowhead indicates a GSC becoming GFP positive. Scale bar: 10µm.

**Movie S2: GFP activation in a GB expressing Bam-FLPD5, nos>>mCD8GFP marking system**

Time-lapse movie of a testis tip showing the GFP-activation in a GB (corresponding to Figure 3D). The testes are expressing Bam-FLPD5, nos>>mCD8GFP marking system, and germline cells are visualized by Vasa-mCherry (magenta). The arrowhead indicates a GB becoming GFP positive. Scale bar: 10µm.

1.	Gadre, P., et al., *The rates of stem cell division determine the cell cycle lengths of its lineage.* iScience, 2021. **24**(11): p. 103232.
